# Transmission and antibiotic resistance of *Achromobacter* in cystic fibrosis

**DOI:** 10.1101/2020.08.04.235952

**Authors:** Migle Gabrielaite, Jennifer A. Bartell, Niels Nørskov-Lauritsen, Tacjana Pressler, Finn C. Nielsen, Helle K. Johansen, Rasmus L. Marvig

## Abstract

*Achromobacter* species are increasingly being detected in patients with cystic fibrosis (CF), and this emerging pathogen is associated with antibiotic resistance and more severe disease outcomes. Nonetheless, little is known about the extent of transmission and antibiotic resistance development in *Achromobacter* infections.

We sequenced the genomes of 101 clinical isolates of *Achromobacter* (*A. xylosoxidans* based on MALDI-TOF/API N20 typing) collected from 51 patients with CF—the largest longitudinal dataset to-date. We performed phylogenetic analysis on the genomes and combined this with epidemiological and antibiotic resistance data to identify patient-to-patient transmission and development of antibiotic resistance.

We found that MALDI-TOF/API N20 was not sufficient for *Achromobacter* species-level typing, and that the population of *Achromobacter* isolates was composed of five different species where *A. xylosoxidans* accounted for 52% of infections. Most patients were infected by unique *Achromobacter* clone types; nonetheless, suspected patient-to-patient transmission cases identified by shared clone types were observed in 35% (N=18) of patients. In 15 of 16 cases the suspected transmissions were further supported by genome- or clinic visit-based epidemiological analysis. Finally, we found that resistance developed over time.

We show that whole-genome sequencing (WGS) is essential for *Achromobacter* species typing and patient-to-patient transmission identification which was identified in *A. ruhlandii, A. xylosoxidans* and, for the first time, *A. insuavis*. Furthermore, we show that the development of antibiotic resistance is associated with chronic *Achromobacter* infections. Our findings emphasize that transmission and antibiotic resistance should be considered in future treatment strategies.

## Introduction

The majority of patients with CF are affected by bacterial airway infections which persist for years and often are the cause of respiratory failure and premature death. (1) *Pseudomonas aeruginosa* remains the most common pathogen causing infections in patients with CF airways (1,2); however, *Achromobacter* is an emerging and less studied opportunistic pathogen. (3,4) Understanding of bacterial antibiotic resistance development and transmission is crucial for effective pathogen management and elimination. (5–8) For example, *A. ruhlandii* Danish Epidemic Strain (DES) has already been defined as a hypermutable and antibiotic-resistant clone type that has been transmitted among Danish patients with CF (9–11).

Here, we sequenced and analyzed the genomes of the largest to-date collection of clinical isolates of *Achromobacter* from patients with CF. First, we aimed to assess how species level *Achromobacter* typing based on WGS compares to species typing based on biochemical or mass spectrometry methods which were used for routine clinical diagnostics. Second, we aimed to use genetic distance and phylogenetic relationship of genomes to identify cases of transmission of *Achromobacter* between patients, and also to include other epidemiological data to identify possible drivers of transmission. Third, we aimed to investigate and present the extent of antibiotic resistance development in the light of genetic epidemiological findings. Overall, we aimed to better understand the patient-to-patient transmission and antibiotic resistance development in *Achromobacter* during infections in patients with CF, ultimately leading to improved strategies to handle persistent airway infections.

## Materials and Methods

### Bacterial isolates

The analysis included 101 clinical isolates of *Achromobacter* that prior to this study were identified in the routine clinical microbiology laboratory as *A. xylosoxidans* by API N20 (bioMérieux, France) or MALDI-TOF typing (Bruker, Germany). The isolates were sampled from 51 patients with CF attending the Copenhagen Cystic Fibrosis Center at Rigshospitalet, Denmark. This data set represents 49% (51 out of 104) of all patients attending Copenhagen Cystic Fibrosis Center with *A. xylosoxidans* detected at least once (as defined by MALDI-TOF or API N20) in years 2002–2018 (detailed description of patients is provided in Table S1). We included isolates sampled before 2002 for nine of the patients; however, samples from patients with *Achromobacter* detected only prior to 2002 were not included in the study. Four isolates were previously analysed by Veschetti *et al*. (2020) (12) where patients A and B correspond to patients P0802 and P8603, respectively, in this study. The use of clinical isolates was approved by the local ethics committee at the Capital Region of Denmark (Region Hovedstaden; approval registration number H-4-2015-FSP), and the use of clinical registry data was approved by the Danish Agency for Patient Safety (approval registration number 31-1521-428).

### Antibiotic treatment

All patients received early antibiotic treatment for *Achromobacter* at the first positive culture. All treatments were based on antibiotic susceptibility testing. The most used treatment regime was inhalations of colistin in combination with amoxicillin-clavulanic acid for 3 weeks. (4) If early eradication treatment failed, other treatment modalities were used; mainly 14 days intravenous treatment with either piperacillin-tazobactam or meropenem, or ceftazidime in combination with tobramycin and sulfatrim. In some cases, patients were treated with inhaled or orally administered colistin or ceftazidime.

### Bacterial genome sequencing and definition of clone type

Genomic DNA was extracted and purified from *Achromobacter* clones with DNeasy Blood and Tissue kit (Qiagen). Genomic DNA libraries were prepared using a Nextera XT DNA Library Prep kit (Illumina), and libraries were sequenced on an Illumina MiSeq instrument generating 250 base paired-end sequencing reads (average of 1,124,551 read pairs; range of 350,677–2,118,817 read pairs). Clone types were defined by Pactyper (13) using the default settings and a species core genome defined by GenAPI using the 101 *Achromobacter* isolates from this study. (14)

### *De novo* assembly-based phylogenetic tree generation

Sequence reads from each isolate were corrected and assembled into scaffolds by SPAdes 3.10.1 (15) using default settings and k-mer sizes ranging from 21 to 127. Genome assemblies consisted on average 216 scaffolded contigs (range 92–506). Core genome SNV-based phylogenetic trees of the 101 *de novo* assembled *Achromobacter* isolates together with publicly available reference genomes were generated with parsnp 1.2 (16) using default settings. Eight complete *Achromobacter* reference genomes were included in the phylogenetic analysis (RefSeq assembly accession: GCF_000165835.1 (*A. aegrifaciens*), GCF_000758265.1 (*A. xylosoxidans*), GCF_001051055.1 (*A. ruhlandii*), GCF_001457475.1 (*A. xylosoxidans*), GCF_001558755.2 (*A. insuavis*), GCF_001558915.1 (*A. ruhlandii*), GCF_001559195.1 (*A. xylosoxidans*) and GCF_900475575.1 (*A. xylosoxidans*)). The phylogenetic tree was visualized with Microreact webservice (17). Phylogenetic tree of *A. ruhlandii* clone type AX01DK01 isolates together with 19 *A. ruhlandii* DES genomes from Ridderberg *et al*. (2020) (10) was based on core genome SNVs using parsnp 1.2 (16) with the default settings and visualized with iTOL webservice (18).

### Patient-to-patient transmission identification

Phylogenies and genetic distances of isolates from patients which shared a clone type and were suspected to participate in patient-to-patient transmission events, were determined with BacDist (to determine SNV-based phylogenetic relationships and pairwise SNV distances) (19) and GenAPI (to determine gene content differences) (14). Phylogenies were visualized using iTOL webservice (18).

In our local CF database, we extracted days and wards at which the patients had registered microbial samples from 2002 through December 2018. The far majority of samples was taken when the patient presented at the ward; nonetheless, we note that a few of the samples were sent to the ward and registered without the patient being present. We used this information to identify days of possible contact between patients (hereafter defined as ‘contact days’), i.e. we were able to infer if patients had potentially been at the same hospital ward on the same date, not if they had actually met at the hospital. Clinic visit dates from two years prior to the first *Achromobacter*-positive sample for each patient recorded in the digital registry were included in the analysis (to account for undetected colonization/transmission) for all 51 patients with sequenced *Achromobacter* in this study. The analysis was performed for each patient pair (1,275 possible combinations).

### Hypermutator identification

Hypermutators were identified in clone types, where two or more isolates were available, by using BacDist (19) to call genetic variants, and then the transition-to-transversion nucleotide substitution ratio (Ts/Tv) was evaluated: if Ts/Tv was >3 the clone type was concluded to be hypermutable. Insertions, deletions and frameshifts in the MMR system *mutL* and *mutS* gene alignments were manually evaluated to identify which genetic changes could cause hypermutability. (20)

### Antibiotic susceptibility testing and statistical analysis

Rosco Diagnostica antibiotic-containing tablets and the corresponding zone of inhibition interpretive breakpoints were used for antibiotic susceptibility testing, and isolate susceptibility profiles were interpreted as ‘Resistant’, ‘Intermediate resistant’, or ‘Susceptible’ according to manufacturer’s guidelines. Antibiotic susceptibility profiles were available for all 92 *Achromobacter* isolates which were sampled since 2002 (Table S2). Antibiotic susceptibility profiles were tested for Amoxicillin-Clavulanate (AMC), Ampicillin (AMP), Aztreonam (ATM), Ceftazidime (CAZ), Ceftriaxone (CRO), Cefuroxime (CXM), Chloramphenicol (CHL), Ciprofloxacin (CIP), Colistin (CST), Imipenem (IPM), Meropenem (MEM), Moxifloxacin (MXF), Penicillin (PEN), Piperacillin-Tazobactam (TZP), Rifampicin (RIF), Sulfamethizole (SMZ), Tetracycline (TET), Tigecycline (TGC), Tobramycin (TOB), Trimethoprim (TMP) and Trimethoprim-Sulfamethoxazole (SXT). Statistical analysis of resistance phenotype distribution differences between early, late and unique isolates was performed using Wilcoxon signed-rank test using 0.05 p-value threshold.

## Results

### Selection of *Achromobacter* isolates for sequencing

From our local CF database, we found that 104 patients attending the Copenhagen Cystic Fibrosis Clinic in years 2002–2018 had at least one positive culture of *Achromobacter* (*Achromobacter xylosoxidans* as defined by MALDI-TOF or API N20). We sequenced the genomes of 101 isolates of *Achromobacter* from 51 of the patients (Figure 1). The isolates from 2002–2015 were primarily selected in order to sequence the first and last isolate from patients showing positive culture of *Achromobacter* over long time periods, and from year 2015 and onwards all isolates from any patient were selected for genome sequencing. Also, we included isolates sampled before 2002 for nine of the patients.

**Figure 1.**
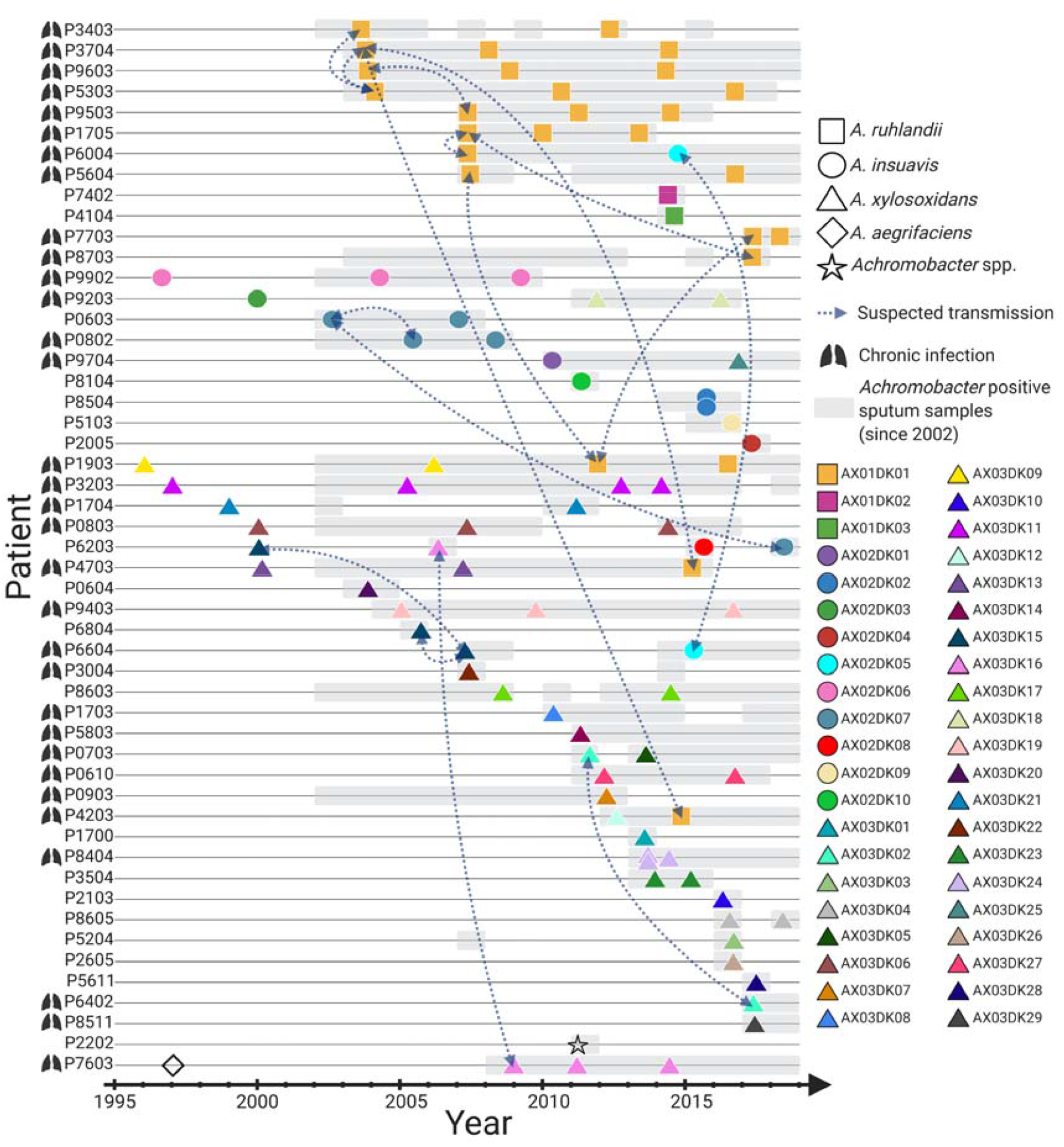
Overview of 101 longitudinally collected *Achromobacter* isolates from patients with CF.

For 30 patients, we sequenced at least two longitudinally collected isolates (range 2–4 isolates), and for 21 patients, we sequenced only a single isolate (N=20) or two isolates from the same time point (N=1; patient P8504). The time between first and last sequenced isolate from patients with longitudinal isolates ranged from 1 to 20 years. Patients P9503 and P9603 were siblings.

Of the 51 patients, 32 (63%) were clinically defined as chronically infected with *Achromobacter*, i.e. the patients had half or more of their samples positive for *Achromobacter* over a year when at least 4 samples were taken, or when specific precipitating antibodies against *Achromobacter* were 4 or more. (21) Of the 53 patients for which no isolates were included in the study, 15 (28%) were clinically defined as chronically infected with *Achromobacter* (Table S1).

### *Achromobacter* species typing

Prior to this study, all included isolates were identified as *A. xylosoxidans* species in the routine clinical microbiology laboratory by MALDI-TOF or API N20 typing.

Nonetheless, when we compared our *Achromobacter* isolate genome sequences to eight publicly available complete *Achromobacter* reference genomes, we found that our isolate collection was composed of five different *Achromobacter* species (Figure 2A). Accordingly, 15 (25%) patients were infected with *A. ruhlandii* (AX01), 12 (20%)—with *A. insuavis* (AX02), 31 (52%)—with *A. xylosoxidans* (AX03), and 2—with other *Achromobacter* species (*A. aegrifaciens* and a new genogroup; AX04). Since only a single isolate was available for *A. aegrifacies* and a new genogroup, respectively, these isolates were excluded from further analysis.

**Figure 2.**
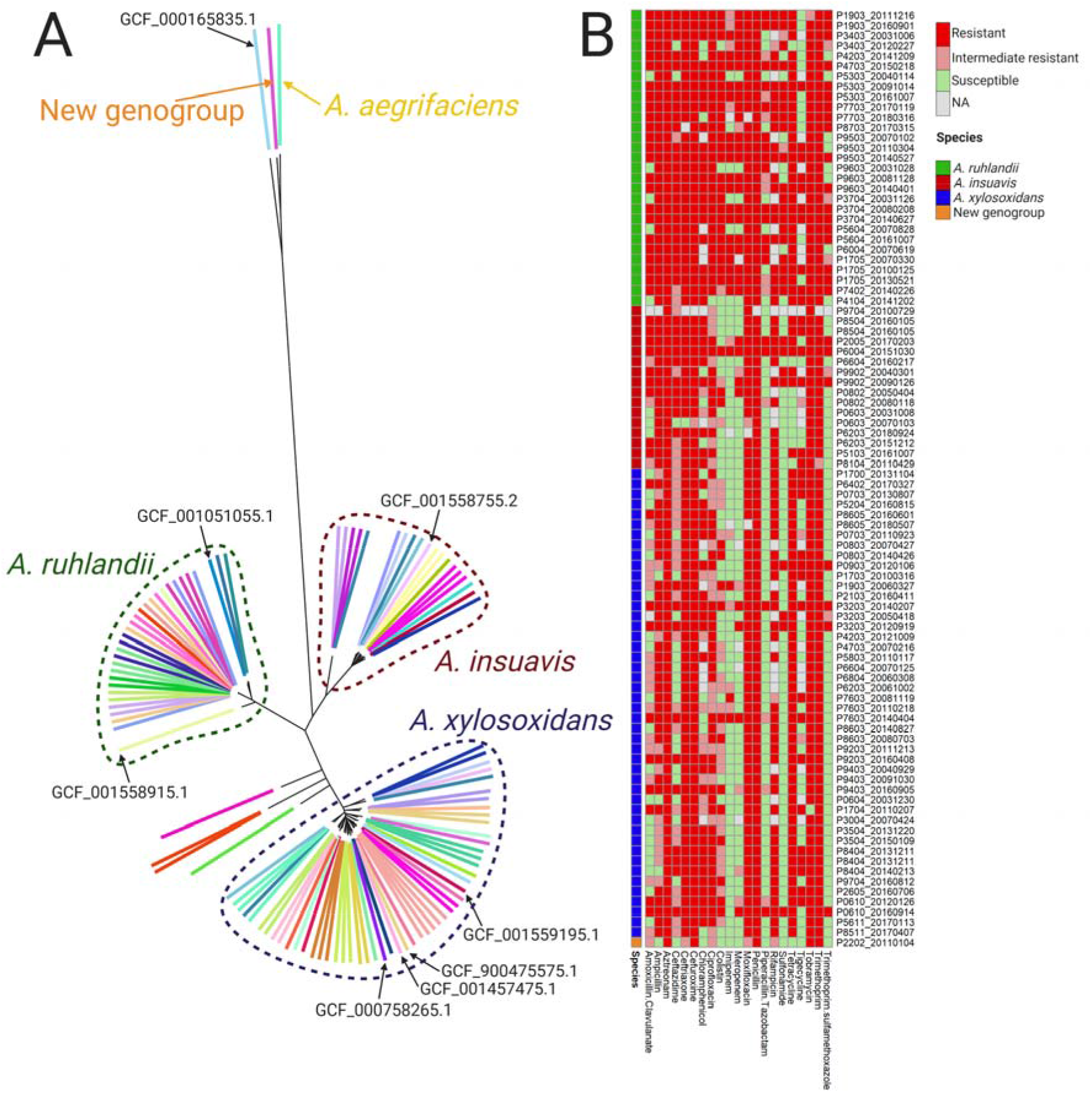
Population structure and susceptibility to antibiotics of *Achromobacter* clinical isolates. (A) Phylogenetic tree of 101 *Achromobacter* clinical isolates together with eight *Achromobacter* reference genomes. Colors represent bacterial isolates from different patients; black arrows point to *Achromobacter* reference genomes. The phylogenetic tree can be accessed on the Microreact webserver (https://microreact.org/project/ByZx4dqC7). (B) Overview of susceptibility profiles of 92 *Achromobacter* isolates against 21 antibiotics.

### Clonal identity of isolates

We further compared the genomes to determine the clonal identity of isolates, i.e. isolates different by less than 5,000 SNVs in the core genome were assigned the same clone type. The minimum observed pairwise distances between isolates from different clone types were 41,237, 32,339 and 8,047 SNVs for *A. ruhlandii, A. insuavis* and *A. xylosoxidans*, respectively, and the maximum observed pairwise distance between isolates belonging to the same clone type were 1,634, 482 and 1,160 SNVs for *A. ruhlandii, A. insuavis* and *A. xylosoxidans*, respectively. Of the 30 patients for which we had longitudinally collected isolates, 21 patients were infected with the same clone type over time, and 10 patients were infected with more than one clone type (N=2) and/or species (N=9) of *Achromobacter* (Figure 1).

### Patient-to-patient transmission in three *Achromobacter* species

Six of the clone types were found in more than one patient (range 2–13); thus, we were interested if sharing of clone types was due to transmission between patients. We identified clonal isolate pairs that represented minimal SNV distance between isolates from different patients, and we defined these pairs as 16 suspected transmission cases for further investigation (Table 1; Figure 3A).

**Table 1.**
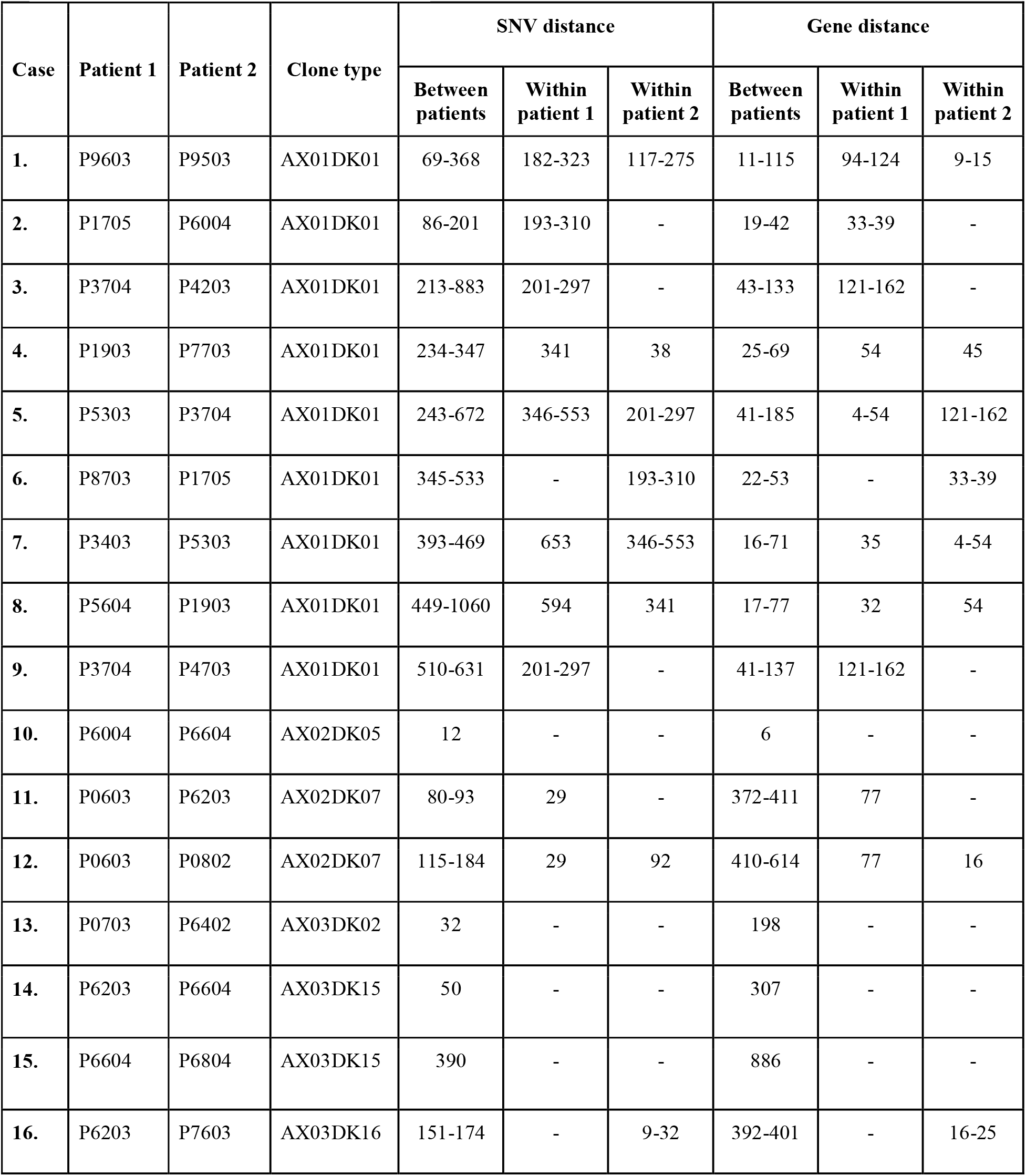
Smallest SNV and gene content distances within- and between-lineages involved in patient-to-patient transmission.

| Case | Patient 1 | Patient 2 | Clone type | SNV distance |  |  | Gene distance |  |  |
| --- | --- | --- | --- | --- | --- | --- | --- | --- | --- |
|  |  |  |  | Between patients | Within patient 1 | Within patient 2 | Between patients | Within patient 1 | Within patient 2 |
| 1. | P9603 | P9503 | AX01DK01 | 69-368 | 182-323 | 117-275 | 11-115 | 94-124 | 9-15 |
| 2. | P1705 | P6004 | AX01DK01 | 86-201 | 193-310 | - | 19-42 | 33-39 | - |
| 3. | P3704 | P4203 | AX01DK01 | 213-883 | 201-297 | - | 43-133 | 121-162 | - |
| 4. | P1903 | P7703 | AX01DK01 | 234-347 | 341 | 38 | 25-69 | 54 | 45 |
| 5. | P5303 | P3704 | AX01DK01 | 243-672 | 346-553 | 201-297 | 41-185 | 4-54 | 121-162 |
| 6. | P8703 | P1705 | AX01DK01 | 345-533 | - | 193-310 | 22-53 | - | 33-39 |
| 7. | P3403 | P5303 | AX01DK01 | 393-469 | 653 | 346-553 | 16-71 | 35 | 4-54 |
| 8. | P5604 | P1903 | AX01DK01 | 449-1060 | 594 | 341 | 17-77 | 32 | 54 |
| 9. | P3704 | P4703 | AX01DK01 | 510-631 | 201-297 | - | 41-137 | 121-162 | - |
| 10. | P6004 | P6604 | AX02DK05 | 12 | - | - | 6 | - | - |
| 11. | P0603 | P6203 | AX02DK07 | 80-93 | 29 | - | 372-411 | 77 | - |
| 12. | P0603 | P0802 | AX02DK07 | 115-184 | 29 | 92 | 410-614 | 77 | 16 |
| 13. | P0703 | P6402 | AX03DK02 | 32 | - | - | 198 | - | - |
| 14. | P6203 | P6604 | AX03DK15 | 50 | - | - | 307 | - | - |
| 15. | P6604 | P6804 | AX03DK15 | 390 | - | - | 886 | - | - |
| 16. | P6203 | P7603 | AX03DK16 | 151-174 | - | 9-32 | 392-401 | - | 16-25 |

**Figure 3.**
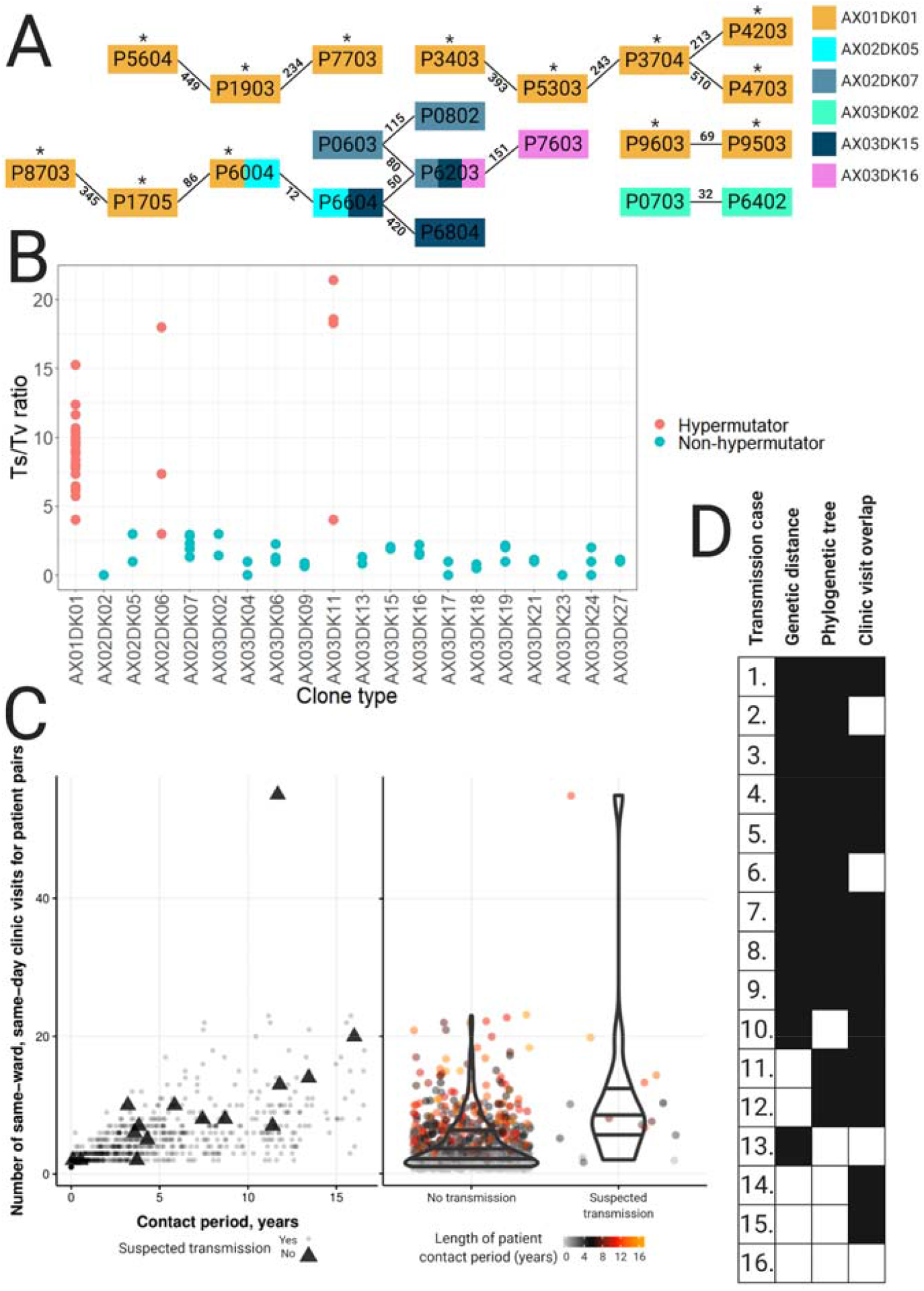
Mutational and transmission analysis of *Achromobacter* isolates. (A) All identified suspected patient-to-patient transmission cases, the smallest pairwise SNV genetic distances between the isolates are stated above the lines. Hypermutators are marked with an asterix. AX01—*A. ruhlandii*, AX02—*A. insuavis* and AX03—*A. xylosoxidans* isolates. (B) Transition-to-transversion substitution ratio for 20 *Achromobacter* clone types. The red horizontal line marks the Ts/Tv ratio of 3. (C) Number of times a patient pair visited the same hospital ward at the same date versus the period of time from first to last potential contact in years. Patients with suspected transmission are marked with triangles (left panel) and a distribution of patient contacts (by number of same-ward, same-day clinic visits) for patient pairs with suspected transmission versus the rest of the patient pair cohort (right panel). Point color corresponds to length of patient contact period in years (from first contact date to last contact date of patient pair). (D) A summary of genetic, phylogenetic and clinic visit overlap support for each suspected transmission case. Instances where cases have support for suspected transmission are colored in black.

In 12 suspected transmission cases (cases 1-9, 11-12, and 16), we had multiple clonal isolates from at least one of the paired patients, enabling us to compare within-versus between-patient genetic diversity (SNV and gene distance). In nine of the 12 cases, we found that isolates from different patients were more closely genetically related than isolates from the same patient (cases 1-9 in Table 1). For example, AX01DK01 isolates from sibling patients P9603 and P9503 differed by 69 SNVs and 8 genes whereas within-patient diversity was 117–323 SNVs and 9–124 genes (Table 1).

Next, we generated phylogenetic trees to reconstruct the genetic relationship of all four suspected patient-to-patient transmitted clone types AX01DK01 (*A. ruhlandii*), AX02DK07 (*A. insuavis*), AX03DK15 (*A. xylosoxidans*), and AX03DK16 (*A. xylosoxidans*) for which three or more isolates were available, to find phylogenetic support for transmission events. Of 14 suspected transmission cases with available phylogenetic information, phylogenetic trees showed support for transmission in 12 of cases (cases 1–9 and cases 11–12 in Table 1) as evidenced by isolates from one patient to be phylogenetic descendants to isolates from another patient (Figure S2).

We did not have enough isolates available for cases 10 and 13 to determine within-patient genetic diversity nor phylogenies. Nonetheless, the clonal isolate pairs in the suspected cases showed only 12 and 32 SNV differences, respectively. As such, the SNV distances between isolates from patients in cases 10 and 13 was less than within-patient genetic distances observed for any of the patients for which we had multiple clonal isolates, except for AX02DK07 isolates from patient P0603 that showed 29 SNV difference. Accordingly, we found that the relatively short genetic distances between patient isolates supported suspicion of transmission for cases 10 and 13.

To add on the genetic evidence of transmission, we analyzed the overlaps of patient visits to the clinic. We used our local CF database to count the number of days, i.e. contact days, at which pairs of patients had microbial sampling at the same hospital ward. Patients were in potential contact with 8–45 other patients between 2002 and 2018. In total, we identified 3,522 patient contact days distributed across 804 patient pairs (out of a possible 1,275 patient pairs). Of the 16 patient pairs with suspected transmission events, only one patient pair (P8703 and P1705) never had microbial sampling in the same hospital ward on the same day. When analyzing all patients with at least one contact day, suspected transmissions tended to happen in patients with more contact days than non-transmission patients (median 8 vs 3 patient contacts, respectively) (Figure 3C). Clinic visit data further supports the suspected transmission in twelve cases (cases 1, 3–5, 7–12, and 14-15; see Table S3 for more information) where potential between-patient contact happened before the first isolation of the transmitted clone type. Overall, 15 out of 16 suspected between-patient transmission cases were supported by genetic distance, phylogenetic and/or epidemiological data (Figure 3D). Only suspected patient-to-patient transmission between P6203 and P7603 had no supporting evidence.

*A. ruhlandii* clone type AX01DK01 showed the most suspected transmissions and was represented by 27 isolates across 13 patients. Therefore, we compared genomes of AX01DK01 isolates to 19 genomes of clone type DES (*A. ruhlandii*), which has previously been reported to be frequently transmitted among Danish CF patients (10). We included genomes of DES isolates from both Copenhagen CF Center (N=12) and Aarhus CF Center (N=7), and our analysis confirmed that clone types AX01DK01 and DES are the same (Figure S1).

### Hypermutators only in chronic infections

Clone type AX01DK01/DES has in a previous report been shown to be hypermutable; putatively due to an in-frame 36-nt deletion in DNA mismatch repair (MMR) gene *mutS* (10). We found the same *mutS* deletion in all AX01DK01/DES isolates from this study, and the isolates showed a large genetic diversity driven by an excess of transition substitutions (Figure 3B; e.g. isolates from patient P3403 were different by 677 SNVs of which 624 were transitions) which is consistent with hypermutation caused by a defective DNA MMR system.

Next, we tested if hypermutation was evident in the other 19 clone types for which two or more isolates were available. We found that AX02DK06 and AX03DK11 of species *A. insuavis* and *A. xylosoxidans*, respectively, also showed excess numbers of transition substitutions; thus, we also defined these two clone types as hypermutable. We also searched for mutations in the DNA MMR genes *mutS* or *mutL* in these two clone types, but we did not find that the identified mutations were always associated with an excess of transition substitutions. Finally, we noted that hypermutable clone types were exclusively found in patients clinically defined as chronically infected (Figure 1).

### Development of antibiotics resistance over time

We were able to retrieve clinical routine diagnostic measurements of susceptibility profiles against 21 antibiotics for all 92 isolates sampled from 2002 and onwards. For the 21 patients where only single isolates were available, the isolates were resistant or intermediately resistant to a median of 14 antibiotics. For the 30 patients for which we included longitudinally collected isolates, we found early and late isolates to be resistant or intermediate resistant to a median of 14 and 18 antibiotics, respectively. Wilcoxon ranked-sum test showed that late isolates were statistically significantly less susceptible than early (p=3.9×10^−3^) and single (p=5.0×10^−4^) isolates. Nearly all (87–92) isolates were resistant or intermediate resistant to these nine antibiotics: ATM, CRO, CXM, CIP, MXF, PEN, RIF, TOB and TMP. In contrast, no antibiotic was effective against all isolates, but many (51–63) isolates were susceptible to these five antibiotics: IPM, MEM, TZP, SMZ and SXT (Figure 2B). Clone type AX01DK01/DES isolates were resistant or intermediate resistant to a median of 20 antibiotics, whereas the median was 14 for other *Achromobacter* isolates. Wilcoxon ranked-sum test showed a significant difference between the two groups (p=2.3×10^−7^). No statistically significant difference between the median number of resistance or intermediate resistance of *A. insuavis* and *A. xylosoxidans* isolates was identified (p=0.92). Interestingly, 7 out of 29 *A. ruhlandii* (6 out of 27 AX01DK01/DES) isolates were resistant or intermediate resistant to all 21 antibiotics while only one *A. insuavis* and none of the *A. xylosoxidans* isolates were resistant or intermediate resistant to all antibiotics.

## Discussion

*Achromobacter* is an emerging pathogen causing chronic respiratory tract infections in patients with CF; however, the genetic epidemiology of these infections is not well understood. We sequenced and analyzed 101 genomes of *Achromobacter* isolates from 51 patients with CF which is the largest longitudinally collected *Achromobacter* genome dataset available to-date.

Our analysis revealed that nearly 20% of patients were infected with 2–4 *Achromobacter* species and/or clone types over the sampled time-period which suggests that not all *Achromobacter* colonizing the airways lead to chronic infections and further supports the early antibiotic treatment of *Achromobacter* infections. (9) Besides, the frequent observation of multiple *Achromobacter* species and/or clone types in the same patient emphasize the necessity for a sensitive species- and clone type-level typing scheme for *Achromobacter* to distinguish infections caused by different *Achromobacter* species and/or different *Achromobacter* clone types. Phylogenetic analysis showed that MALDI-TOF/API 20 is not accurate for *Achromobacter* species-level typing as it has also been recently indicated by others (22–24); thus, we suggest that sequencing of marker genes (e.g., *bla*_*OXA*_ or *nrdA*) or WGS should be used for species-level typing in the clinical setup. (25) WGS, furthermore, can facilitate patient-to-patient transmission identification.

The majority of *Achromobacter* infections are acquired from the environment and prevalence of patient-to-patient transmission remains controversial: while some studies identified patients being infected with unique clone types of *Achromobacter* (26–28) other studies reported the cases of suspected patient-to-patient transmission. (7,21,29,30) We suspected cases of transmission between patients for all three *Achromobacter* species in our study. Of 16 suspected transmission cases, 15 were supported by genetic distance measurements, phylogenetic trees, and/or epidemiological data. Transmission between P1705 and P6004 patients was previously defined as an indirect transmission event by Hansen *et al*. (2013). (21) P9603 and P9503 are siblings who share the same living environment; therefore, transmission of CF airway pathogen between them is highly likely. Between-patient transmission of *A. insuavis* has not been described in the scientific literature before.

We note that the observed SNV distance between the transmitted *Achromobacter* isolates is higher than what has been observed for transmitted *P. aeruginosa* (31); hence, this complicates the identification of transmission and might explain why no patient-to-patient transmission was discovered in several previous studies. (26–28) We furthermore showed that gene content differences between isolates could serve as an additional criterium for between-patient transmission identification.

Interestingly, AX01DK01/DES (*A. ruhlandii*)—a known hypermutator—is widely transmitted between patients with CF within and between CF centers in Copenhagen and Aarhus even though some studies show reduced transmissibility of hypermutator strains. (32) We showed that isolates did not phylogenetically group according to center origin, suggesting multiple transmission events between centers. This phenomenon might be explained by patients with CF physically moving from one city and CF center to another. Furthermore, recent first-time isolation of AX01DK01/DES in two patients with CF (P7703 and P8703) indicates that, while transmission of AX01DK01/DES has previously been recognized (10), additional actions might need to be taken to prevent transmission of AX01DK01/DES.

Finally, we observed that infection of the same clone type in several patients was generally a sign of suspected patient-to-patient transmission of *Achromobacter*. This knowledge could be applied in diagnostics where sharing an *Achromobacter* clone type would be the first sign of suspected transmission between patients and would lead to further investigations.

Moreover, we confirmed previous findings that *A. ruhlandii* clone type AX01DK01/DES is hypermutable, and in addition we found evidence of hypermutation in *A. insuavis* and *A. xylosoxidans* clone types. (10) Hypermutation could be pivotal for *Achromobacter* persistence since all hypermutable clone types caused chronic infections in patients with CF. We confirmed that *Achromobacter* is highly antibiotic resistant and furthermore that late isolates from patients which were colonized with *Achromobacter* were significantly more antibiotic resistant than early or single isolates. This indicates that *Achromobacter* adapts to the host environment where high levels of antibiotics are present even though *Achromobacter* is innately highly antibiotic resistant. Moreover, significantly higher antibiotic resistance of AX01DK01/DES than other *Achromobacter* isolates urges for additional efforts to prevent AX01DK01/DES transmission between patients with CF.

Our study has several limitations. First, while our dataset spans over several decades, the number of isolates for each patient is low and we have selected for isolates to represent patients with longitudinal samples leading to overrepresentation of chronically infected patients. More sequenced isolates would have allowed to more accurately identify suspected transmission events, and to better follow hypermutability and antibiotic resistance development. Furthermore, we used single isolates to represent a heterogeneous *Achromobacter* population in patients with CF which could have led to the lack of evidence in some suspected between-patient transmissions. Finally, we did not know if patients have met outside the hospital or during microbial sampling, and we used only potential contact information which adds to the uncertainty about patient contacts. Nevertheless, our study provides evidence for between-patient transmission in all three *Achromobacter* species, development of hypermutability caused by deletions in *mutS* gene, and development of antibiotic resistance over time.

## Conclusions

In summary, by genome sequencing the largest dataset to-date of *Achromobacter* clinical isolates from patients with CF, we showed that MALDI-TOF or API N20 are unsuitable for *Achromobacter* species-level typing, and we conclude that WGS is the most appropriate for species-level typing and patient-to-patient transmission identification. Moreover, we confirmed that sequencing of only one isolate is not sufficient as multiple patients were infected with different species or clone types of *Achromobacter*. For the first time we identified suspected between-patient transmission in *A. insuavis*. Furthermore, we found genomic and epidemiological support for suspected patient-to-patient transmission in all three *Achromobacter* species and suggest a new measure—gene content difference—to be taken into account when evaluating suspected between-patient transmission cases. We, furthermore, emphasized that a shared clone type is the first sign of possible patient-to-patient transmission. Finally, we showed that antibiotic resistance develops in all three *Achromobacter* species and our analysis confirmed previous findings that hypermutability is associated with chronic *Achromobacter* infections. The results of this work allow us to better understand antibiotic resistance dynamics and patient-to-patient transmission of *Achromobacter* in patients with CF which could help predict clinical progression of *Achromobacter* infections and prevent patient-to-patient transmission.

## List of abbreviations

AMC: Amoxicillin-Clavulanate
AMP: Ampicillin
API N20: Analytical profile index N20
ATM: Aztreonam
CAZ: Ceftazidime
CRO: Ceftriaxone
CXM: Cefuroxime
CHL: Chloramphenicol
CIP: Ciprofloxacin
COG: Clusters of Orthologous Groups
CST: Colistin
CF: Cystic fibrosis
DES: Danish epidemic strain (*A. ruhlandii*)
IPM: Imipenem
MALDI-TOF: Matrix assisted laser desorption ionization-time of flight mass spectrometry
MEM: Meropenem
MMR: Mismatch repair
MRCA: Most recent common ancestor
MXF: Moxifloxacin
PEN: Penicillin
TZP: Piperacillin-Tazobactam
RIF: Rifampicin
SMZ: Sulfamethizole
SNV: Single nucleotide variant
TET: Tetracycline
TGC: Tigecycline
TOB: Tobramycin
Ts/Tv: Transition-to-transversion ratio
TMP: Trimethoprim
SXT: Trimethoprim-Sulfamethoxazole
WGS: Whole genome sequencing

## Availability of data and materials

*Achromobacter* whole genome sequencing data is available at European Nucleotide Archive under study accession number PRJEB39108.

## Acknowledgments

Ulla Johansen is thanked for expert technical assistance and Niels Høiby is thanked for collecting the earliest *Achromobacter* isolates. All figures were partly or completely created using BioRender (https://biorender.com/).

## Funding

This work was supported by the Danish Cystic Fibrosis Association (Cystisk Fibrose Foreningen) and the Danish National Research Foundation (grant number 126). HKJ was supported by The Novo Nordisk Foundation as a clinical research stipend (NNF12OC1015920), by Rigshospitalets Rammebevilling 2015-17 (R88-A3537), by Lundbeckfonden (R167-2013-15229), by Novo Nordisk Fonden (NNF15OC0017444 and NNF19OC0056411), by RegionH Rammebevilling (R144-A5287) by Independent Research Fund Denmark / Medical and Health Sciences (FTP-4183-00051) and by ‘Savværksejer Jeppe Juhl og Hustru Ovita Juhls mindelegat’.

## Disclosure declaration

We declare no conflict of interest.

## Ethics declarations

Use of the stored clinical isolates was approved by the local ethics committee at the Capital Region of Denmark RegionH (registration number H-4-2015-FSP).

## Supplementary information

**Table S1**. List of all patients attending Copenhagen CF Center who were infected with *Achromobacter* in 2002–2018.

**Table S2**. Isolate susceptibility data to 21 tested antibiotics. R—resistant to the antibiotic, I— intermediately resistant to the antibiotic, S—susceptible to the antibiotic.

**Table S3**. Suspected transmission cases with clinic visitation overlap information and comments whether the epidemiological data supports the transmission.

**Figure S1**. Core genome SNV-based phylogenetic tree from *de novo* assembled DES isolates of which 7 were from patients attending CF center in Aarhus (blue label background; (10)) and 25 were from patients attending CF center in Copenhagen. The four patient clusters are separated by branch and label colors. The phylogenetic tree can be accessed on iTOL webserver: https://itol.embl.de/tree/12807362379081584977362.

**Figure S2**. Core genome SNV phylogenetic trees of suspected bacterial isolates transmission between patients in *Achromobacter ruhlandii, A. insuavis*, and *A. xylosoxidans* where 4 or more isolates were available. Isolates are named as follows: [Patient ID] [Sampling date] [Species and clone type].

## Notes

### Competing Interest Statement

The authors have declared no competing interest.

