## Supplementary figures and images for "Transmission and antibiotic resistance of *Achromobacter* in cystic fibrosis"

### Figure S1

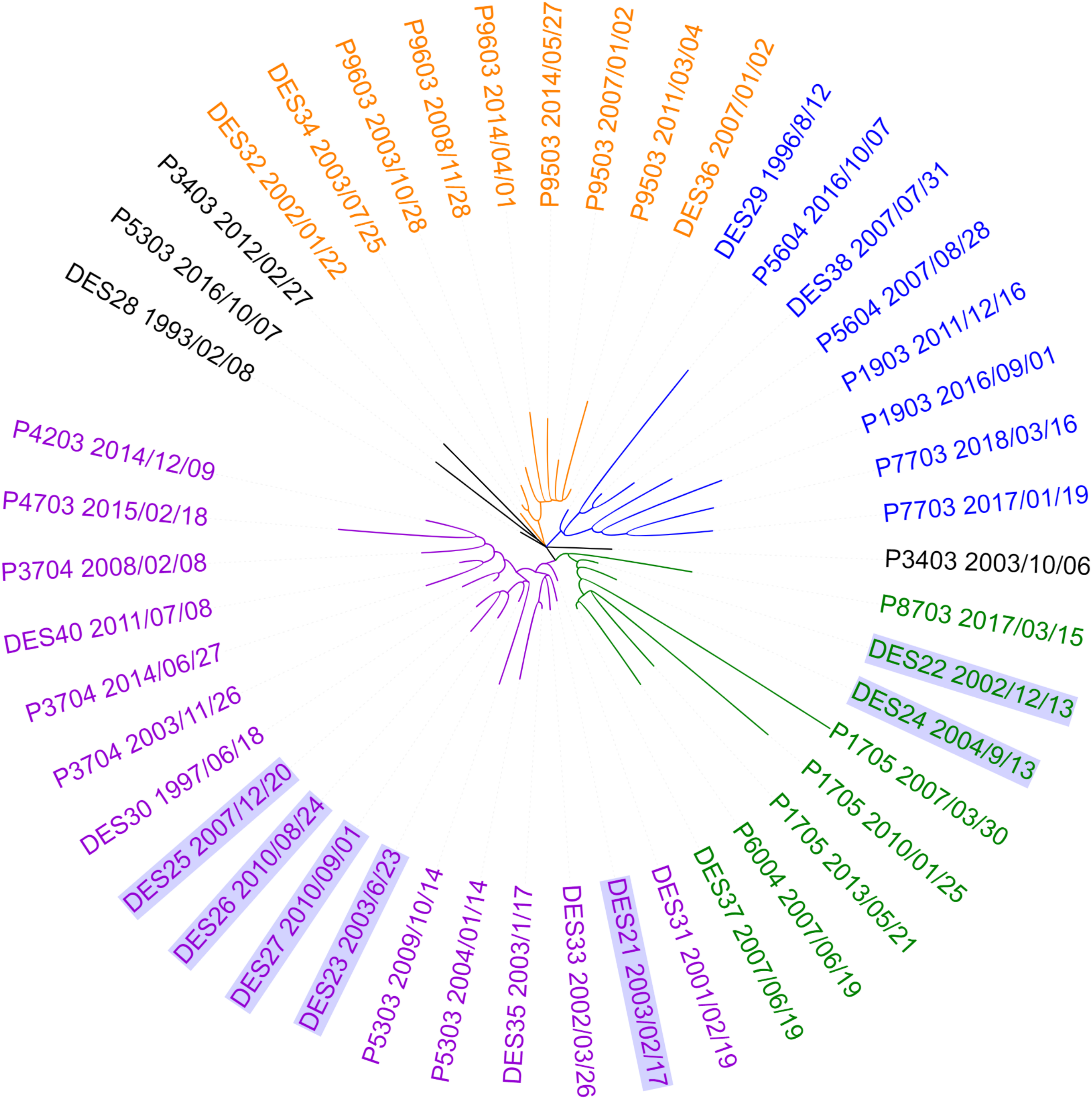

### Figure S2

#### A. ruhlandii

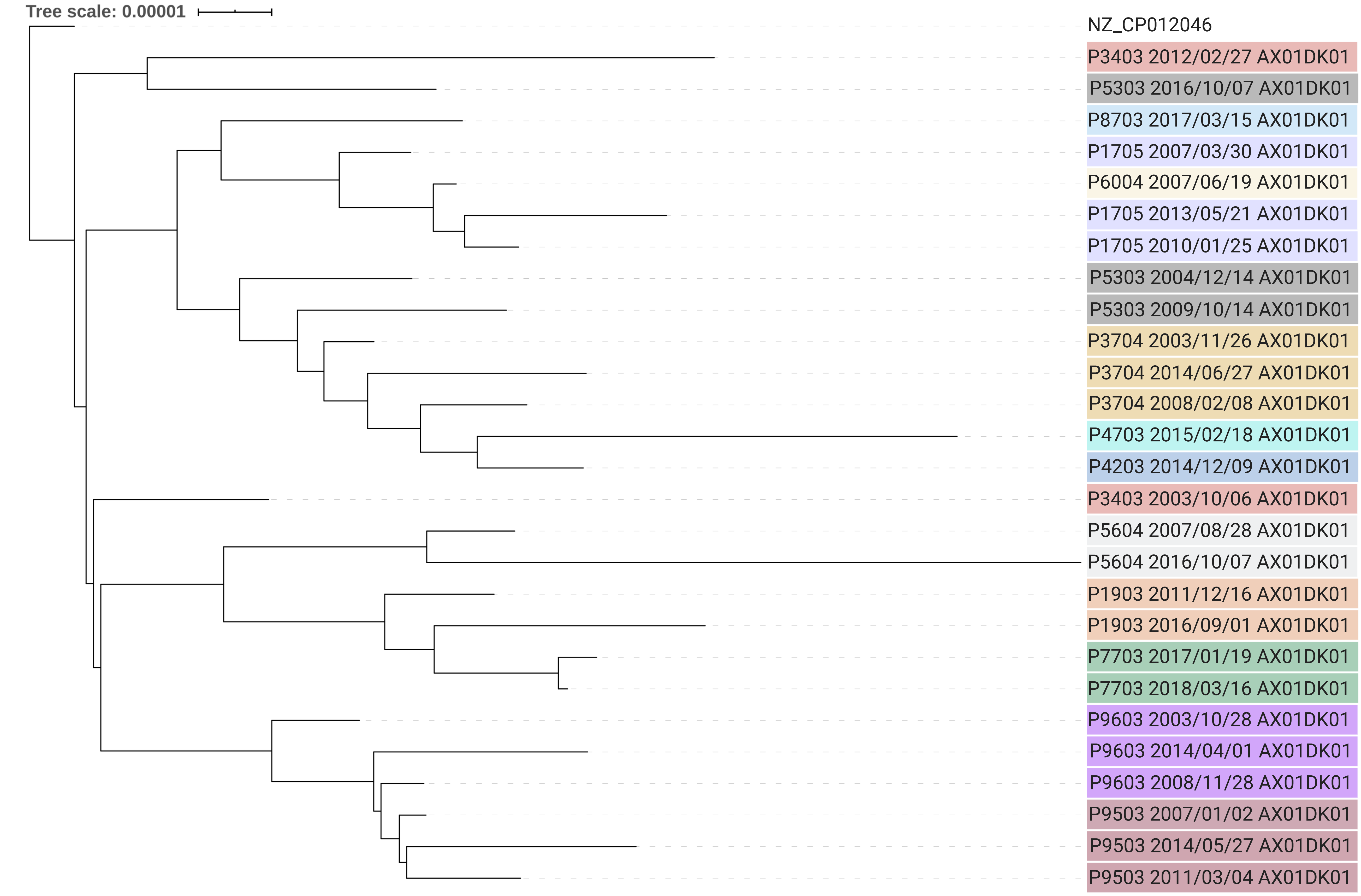


#### A. insuavis

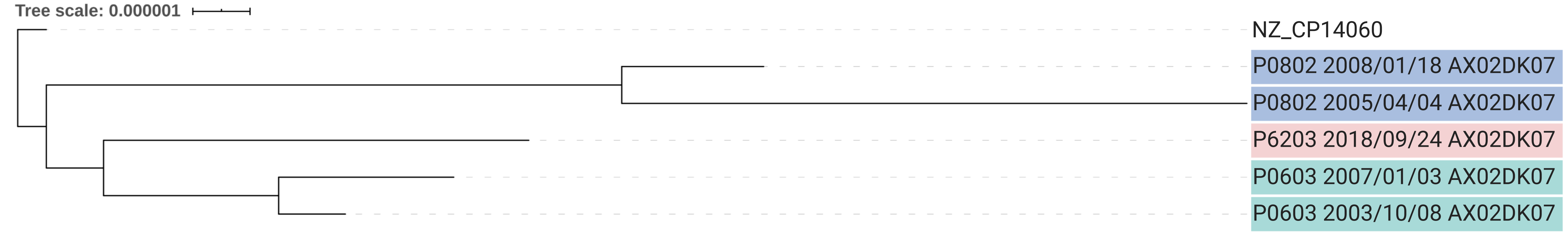


#### A. xylosoxidans

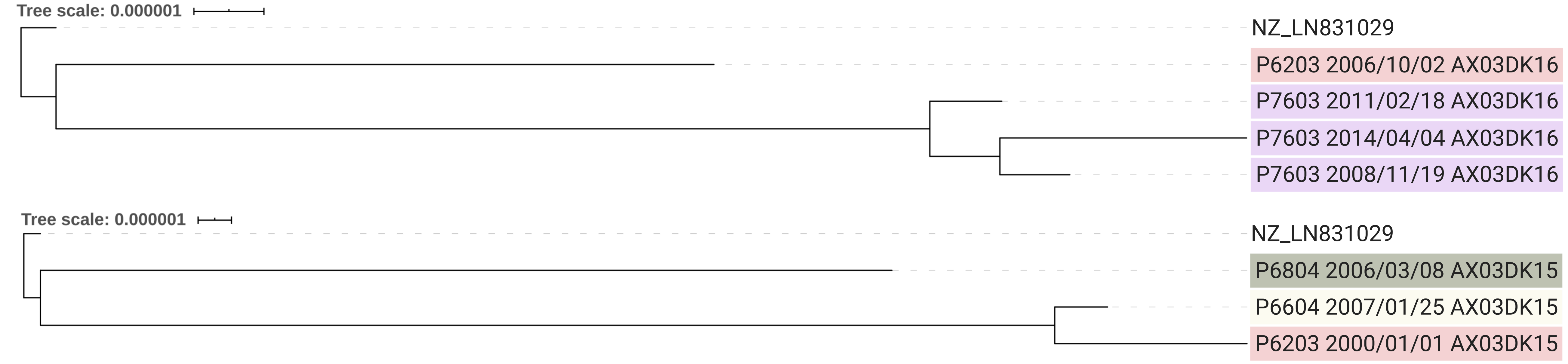
